# Mon1a and FCHO2 are required for maintenance of Golgi architecture

**DOI:** 10.1101/2023.07.06.547837

**Authors:** Dustin C. Bagley, Scott G. Morham, Jerry Kaplan, Diane M. Ward

**Affiliations:** Department of Pathology, Division of Microbiology and Immunology School of Medicine, University of Utah, Salt Lake City Utah, 84132; MesaGen, LLC BioInnovations Gateway, Salt Lake City Utah, 84115

**Keywords:** Golgi, secretory pathway, mitosis, Mon1a, FCHO2

## Abstract

Mon1a has been shown to function in the endolysosomal pathway functioning in the Mon1-Ccz1 complex and it also acts in the secretory pathway where it interacts with dynein and affects ER to Golgi traffic. Here we show that Mon1a is also required for maintenance of the Golgi apparatus. We identified the F-BAR protein FCHO2 as a Mon1a-interacting protein by both yeast two-hybrid analysis and co-immunoprecipitation. siRNA-dependent reductions in Mon1a or FCHO2 resulted in Golgi fragmentation. Membrane trafficking through the secretory apparatus in FCHO2-depleted cells was unaltered, however, reduction of FCHO2 affected the uniform distribution of Golgi enzymes necessary for carbohydrate modification. Fluorescence recovery after photobleaching analysis showed that the Golgi ministacks in Mon1a- or FCHO2-silenced cells did not exchange resident membrane proteins. The effect of FCHO2 silencing on Golgi structure was partially cell cycle-dependent and required mitosis-dependent Golgi fragmentation, whereas the effect of Mon1a-silencing on Golgi disruption was not cell cycle-dependent. mCherry-FCHO2 transiently colocalized on Golgi structures independent of Mon1a. These findings suggest that Mon1a has functions throughout the secretory pathway including interacting with dynein at the ER-Golgi interface in vesicle formation and then interacting with FCHO2 at the Golgi to generate lateral links between ministacks, thus creating Golgi ribbons.

## 1. Introduction

Mon1 is an evolutionarily conserved protein involved in membrane trafficking (1–4). Many organisms have a single Mon1 which affects the endocytic pathway where it acts with Ccz1 as a GEF for Ypt7 (5–8) and is required for all fusion events involving the vacuole including autophagy, cytoplasm to vacuole targeting (Cvt) and multivesicular body (MVB) pathways (6, 9, 10). In the nematode *Caenorhabditis elegans* and in Drosophila melanogaster, Mon1 is required for Rab7 activity, which is necessary for the transition from early to late endosomes as well as secretory granule maturation (4, 11, 12). *C. elegans-*Mon1, SAND1, is thought to displace the activator of Rab5, Rabex5, allowing for the recruitment of Rab7 to endosomal surfaces, thus permitting maturation to a late endosomal compartment (3, 11).

Vertebrates have two homologues of Mon1, Mon1a and Mon1b, which share less than 50% similarity at the amino acid level. Reduction in Mon1a or Mon1b levels alone had no reported effect on endosomal morphology or maturation, however, reductions in both Mon1a and Mon1b showed an increase in the size of the Rab5 compartment and a concomitant decrease in the Rab7 endosomal compartment, suggesting an effect on endosomal maturation (3). The Mon1-Ccz1 complex in mammals has been confirmed to activate Rab7 (13). Further studies have shown a role for Mon1 and the Mon1-Ccz1 complex in secretory lysosomes in osteoclasts (14). These observations suggest that in vertebrates the Mon1 proteins may have expanded roles for other membrane trafficking events. Indeed, the Mon1 protein and the Mon1-Ccz1 complex shows many interacting partners that are involved in membrane trafficking events (14–16). (For review see (7)).

Mon1a was originally identified as modifier of the trafficking of cell-surface and secreted molecules (17). Silencing studies demonstrated that RNAi-mediated depletion of Mon1a resulted in decreased expression of plasma membrane proteins and decreased protein secretion but had no discernable effect on endocytosis (17). Further studies demonstrated that Mon1a played a role in ER to Golgi traffic, as a reduction in Mon1a decreased ER vesicle formation. Protein interaction studies found that Mon1a associated with the microtubule (MT)-based motor dynein and RNAi-mediated loss of either protein reduced ER to Golgi traffic (1).

Evidence that Mon1a acts at the ER does not preclude it from having roles in other trafficking steps. Notably, reductions in Mon1a showed a mild morphological effect on the Golgi apparatus (1). Here we show that varying the levels of Mon1a by siRNA has a significant effect on Golgi morphology and that Mon1a interacts with FCHO2, a protein involved in membrane bending and curvature stabilization at the plasma membrane. Our data suggest that FCHO2 is a Mon1a-interacting protein required for Golgi ribbon formation following mitosis.

## 2. Methods

### 2.1 Mammalian cells and constructs

NIH3T3, Hela, and GalNAc-T2-GFP expressing Hela cells were maintained in DMEM with 10% FBS plus or minus 0.4 g/L G418 (Sigma-Aldrich, St. Louis, MO). GalNAc-T2-GFP Hela cells were a generous gift from Dr. Brian Storrie (University of Arkansas). VSVGtsGFP were a generous gift from Dr. Jennifer Lippincott-Schwartz (National Institutes of Health). Mouse Mon1a was cloned into pCMV-Tag2BFLAG (Stratagene, La Jolla, CA) or pEGFPC1 (Clontech, Mountain View, CA). Mouse Fcho2 was cloned into pCMV-Tag3B-myc or pmCherry-C1. Human FCHO1 was cloned into pEGFPC1 (Clontech, Mountain View, CA). All constructs were sequence verified prior to use.

### 2.2 Transfections and Western analysis

HeLa, NIH3T3 or GalNAc-T2-GFP cells were plated onto tissue culture plates and allowed to grow for 24 to 48 h to 50-80% confluence. The cells were transfected with various constructs using Amaxa nucleofector technology (Lonza, Walkersville, MD) according to the manufacturer’s directions. Protein expression was determined by solubilizing 2–4 x 10^6^ cells in lysis buffer (1% Triton X-100, 150 mM NaCl, 0.5 mM EDTA) plus 2X protease inhibitor cocktail (Roche Applied Science, Boulder, CO). Samples were analyzed by SDS-PAGE and Western analysis was performed using either mouse anti-FLAG antibody (1:1000; Sigma-Aldrich, St. Louis, MO); rabbit anti-GFP (1:5000, ab6556; Abcam); mouse anti-tubulin (1:1000; GeneTex, San Antonio, TX), rabbit anti-FCHO1 (1:500, Abcam ab84740) and mouse anti-Dyn-IC (1:2000, MMS-400R Covance, Princeton, NJ) followed by either peroxidase-conjugated goat anti-mouse immunoglobulin IgG (1:10,000; Jackson ImmunoResearch Laboratories, West Grove, PA), peroxidase-conjugated goat anti-rabbit IgG (1:10,000; Jackson ImmunoResearch Labs, West Grove, PA) or rabbit TrueBlot HRP conjugate anti rabbit IgG 18-8816-33 (eBioscience, San Diego, CA). Antibodies to Mon1a and mouse Fcho2 were generated as described (1) and against the mouse peptide EPKRYRIEIKPAHP (aa327-340 of Fcho2). Rabbit anti-Mon1a and Fcho2 were used at a concentration of 1:500 followed by peroxidase conjugated goat anti-rabbit IgG (1:10,000) (Jackson ImmunoResearch Labs, West Grove, PA). The blots were developed using Western Lightning reagent (PerkinElmer Life Sciences, Waltham, MA). Tubulin was used as a loading control. All experiments were performed a minimum of three times and error bars represent the standard error of the mean.

### 2.3 Size exclusion chromatography

Cells expressing FLAG-tagged Mon1a were lysed in TBS pH 7.4 containing 0.1% Triton X-100 at 0°C for 45 minutes. The lysate was spun down at 14,000 RPM for 10 minutes and filtered through a 0.2-micron filter before running over a FPLC HiLoad 16/20 Superdex 200 prep grade column (Amersham Biosciences, Pittsburg, PA) at 0.5 mL/minute collecting 1 mL fractions. The column was normalized using molecular weight markers including Blue Dextran, thyroglobulin, apoferritin, IgG, bovine serum albumin, horseradish peroxidase and cytochrome C (Sigma-Aldrich, St. Louis, MO). The presence of FLAG-Mon1a in the FPLC fractions was resolved by SDS-PAGE and Western blot analysis.

### 2.4 Yeast two-hybrid (Y2H) and co-immunoprecipitation protein interaction studies

A Mon1a protein fragment containing the first 200 amino acid residues was used as a bait to screen macrophage, spleen and brain libraries for potential interacting partners in budding yeast as described previously (18). Cells were transfected with pFLAG-Mon1a or GFP-Mon1a, solubilized in lysis buffer, incubated 0°C, 30 min, centrifuged at 10,000 x g, 10 min, immunoprecipitated using mouse anti-FLAG antibody (Sigma, St. Louis, MO) or rabbit anti-GFP (GeneTex, San Antonio, TX) and protein A/G-plus agarose (Santa Cruz Biotech, Santa Cruz, CA). Proteins in the immunoprecipitate were identified using Western blot analysis as described above (1).

### 2.5 Treatment with siRNA oligonucleotide pools

Cells were treated with nonspecific (NS), FCHo2, FCHo1, Rab6a or dynein heavy chain1-specific oligonucleotides (Dharmacon SiGenome SiRNA SMARTpool; Dharmacon RNA Technologies, Layfette, CO and Mon1a (5’-UTR)(sense-5’GACUAGGAGCCCAGAACCdt3’ and antisense-5’UGGUUCUGGGUCCUAGUCdt3’), Mon1a (ORF)(sense-5’CUACUACAGCGUUGCCCAAdt3’ and antisense-5’UUGGGCAACGCUGUAGUAGdt3’), University of Utah DNA/RNA synthesis Core; Salt Lake City, UT) using Oligofectamine Reagent (Invitrogen, Carlsbad, CA) according to the manufacturer’s instructions as previously described (1). Briefly, 150,000 cells were treated with siRNAs in OptiMEM reduced-serum medium (Invitrogen, Carlsbad, CA) for six hours at 37°C before serum-replete medium was added for overnight growth. Cells were allowed to grow in DMEM with 10% FBS for 48 or 72 hours before analysis were completed.

### 2.6 Brefeldin A treatment

Cells plated on glass coverslips were incubated with 5 µg/mL Brefeldin A (BFA) (Epicentre Biotechnologies, Madison, WI) for 30 min, washed three times and placed in growth media for recovery. BFA recovery was visualized using an Olympus BX51 upright microscope with a 100X 1.4NA objective and Pictureframer software (Olympus, Melville, NY).

### 2.7 Epifluorescence and Electron Microscopy

Confocal images were captured on a Nikon A1R with the 488nm laser line and a 60x PLANAPO Oil immersion objective. Image analysis was performed using Volocity software as previously described (1). For electron microscopy (EM), cells were fixed in 2.5% glutaraldehyde/1% paraformaldehyde and transmission EM images capture at the University of Utah EM Core as previously described (1).

### 2.8 FRAP analysis

Cell transfected with siRNAs for nonspecific or FCHo2 were analyzed for lateral diffusions and membrane fusion of Golgi GalNAc-T2-GFP by photobleaching and recovery. FRAP was performed using a Nikon A1R with the 488nm laser line and a 60X PLANAPO Oil immersion objective. Regions of interest were selected for photobleaching and after three initial frames, 100% laser power was applied to these regions for two seconds, followed by time-lapse imaging of the recovery at one frame a second for five minutes. Data for the first two minutes were used to generate plots.

### 2.9 Other procedures

Alexa594 conjugated GSII lectin (Invitrogen, Carlsbad, CA) staining was performed as described (19). Protein determinations were performed using bicinchoninic acid protein assay (ThermoFisher).

### 2.10 Statistical Analysis

Data are expressed as mean ± SD from at least three independent experiments unless stated otherwise. Statistical analyses were performed using student’s *t* test with GraphPad Prism software. Outliers were identified in Prism using ROUT (Q=1.0%). P value of less than 0.05 was considered statistically significant and was indicated by asterisks; * p ≤ 0.05, ** p ≤ 0.01, ***, p ≤ 0.001, ****, p ≤ 0.0001.

## 3. Results

### 3.1 Mon1a interacts with the F-BAR protein FCHO2

We demonstrated that Mon1a is required for efficient anterograde trafficking in the secretory pathway due to its interaction with the microtubule-based molecular motor dynein (1). A yeast two-hybrid (Y2H) screen was performed to identify additional interacting partners of Mon1a. A protein fragment containing the first 200 amino acid residues of Mon1a, used as bait, was found to interact with FCHO2 in three independent libraries (**Table 1**). This Mon1a construct also interacted with other proteins but only in single libraries, therefore, we focused on FCHO2 as a binding partner. FCHO2 is an F-BAR domain-containing protein that functions at the cell-surface as part of the machinery necessary for clathrin-dependent endocytosis (20). The yeast two-hybrid result was corroborated by size exclusion chromatography and coimmunoprecipitation. Lysates from cells over-expressing functional FLAG-tagged Mon1a protein (1) were passed over a size-exclusion column. Mon1a is predicted to be a 62 kDa protein, yet the majority of Mon1a was found in two distinct sets of fractions, a ∼150 kDa fraction and in the void volume, which contains molecules greater than 330 kDa (**Figure 1A**). These results suggest that Mon1a acts in two complexes. We hypothesized that these complexes may contain more than one copy of Mon1a. To address this possibility FLAG- and GFP-tagged Mon1a were co-expressed in NIH3T3 cells, GFP-Mon1a was immunoprecipitated and the presence of FLAG-tagged Mon1a assessed by Western blot. Immunoprecipitated GFP-Mon1a was able to co-precipitate FLAG-Mon1a (**Figure 1B**). This interaction was specific as free GFP was unable to co-immunoprecipitate FLAG-Mon1a. Although the transfection of FLAG-Mon1a was weak, as shown in the lysates, FLAG-Mon1a consistently coimmunoprecipitated with GFP-Mon1a supporting the hypothesis that Mon1a complexes have more than one copy of Mon1a.

**Figure 1.**
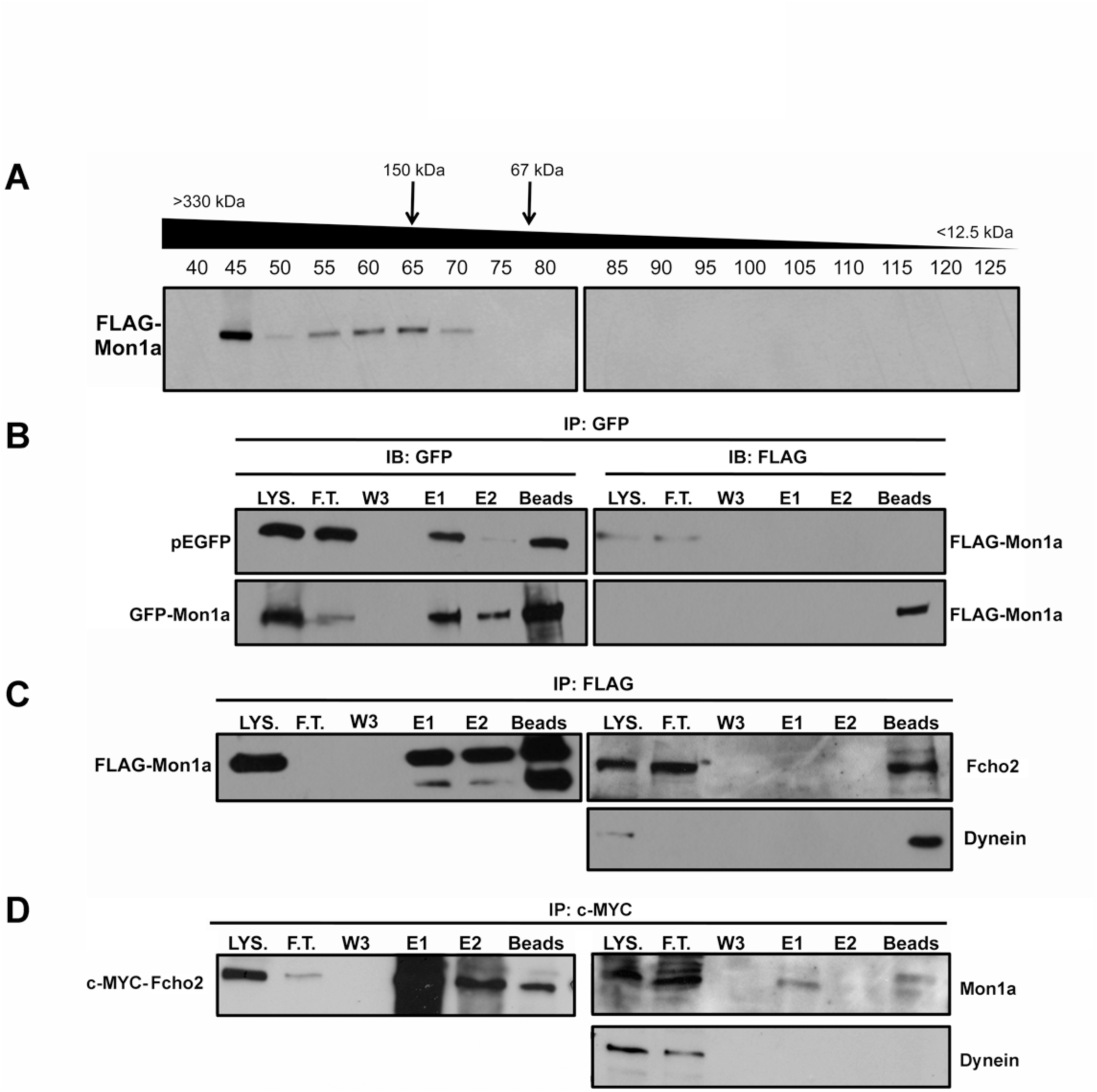
Mon1a associates with a complex containing the endocytic F-BAR domain protein Fcho2. A. NIH3T3 cells were transfected by with FLAG-Mon1a before lysing in the presence of 0.1% Triton X-100. Whole cell lysates were passed through a filter before injecting over a FPLC size-exclusion column. Every 5^th^ fraction between 45-125 was run on SDS-PAGE and Western analysis performed using mouse anti-FLAG antibodies. B. pEGFP, GFP- and FLAG-Mon1a constructs were co-expressed in NIH3T3 cells and lysed. Lysates were incubated with a GFP antibody and A/G-plus sepharose beads, washed extensively, eluted in 0.1M glycine pH 2.5 and samples were resolved by SDS-PAGE and Western blot analysis for co-immunoprecipitation. LYS=lysate (5% loaded), FT=flow through (5% loaded), W3=wash 3, E1=elution one and E2=elution two (20% loaded), beads (50% loaded). C. Cells expressing FLAG-Mon1a were lysed and incubated with FLAG antibody and A/G-plus beads to immunoprecipitate FLAG-Mon1a. SDS-PAGE and Western analysis were done for FLAG, endogenous Fcho2 and dynein. D. Cells expressing c-Myc-Fcho2 were lysed and incubated with anti-c-Myc agarose beads to immunoprecipitate c-Myc-Fcho2. SDS-PAGE and Western analysis were done to confirm c-Myc-Fcho2 immunoprecipitation and probed for endogenous Mon1a and dynein coprecipitations.

**Table 1.** Yeast two-hybrid (Y2H) analysis of Mon1a binding partners. A yeast two hybrid screen was performed using human Mon1a as bait. The first 200 amino acids of Mon1a were shown to interact with the clathrin-dependent endocytosis effector FCHO2. This interaction was seen in multiple independent libraries constructed from macrophages, spleen and brain tissues. Mon1a was also shown to physically associate with Dpy30-like protein (99), a protein thought to be involved in the methylation of histones, tyrosine-protein kinases Fyn and Lyn, thought to be involved in several biological processes including cell growth and activation, respectively. As well as thioredoxin (Txn), which is required for cellular redox signaling.

| **Bait** | **Amino acids** | **Prey** | **a.a. cords.** | **Library** |
| --- | --- | --- | --- | --- |
| Mon1a (*Hs*) | 1-200 | FCHO2 | 13-177 | Macrophage |
| Mon1a (*Hs*) | 1-200 | FCHO2 | 7-106 | Spleen |
| Mon1a (*Hs*) | 1-200 | FCHO2 | 14-150 | Brain |
| Mon1a (*Hs*) | 1-250 | DPY30(99) | 36-100 | Macrophage |
| Mon1a (*Hs*) | 1-200 | FYN(537) | 97-219 | Brain |
| Mon1a (*Hs*) | 1-200 | LYN(582) | 78-276 | Spleen |
| Mon1a (*Hs*) | 1-200 | TXN | -18-106 | Spleen |

To address whether Mon1a interacts with Fcho2, FLAG-Mon1a was immunoprecipitated from the high molecular weight FPLC fraction and the immunoprecipitate examined for the presence of Fcho2. FLAG-Mon1a coprecipitated endogenous Fcho2 as well as dynein, a protein known to interact with Mon1a (1) (**Figure 1C**). The co-precipitation of dynein was stronger than the co-precipitation of Fcho2 suggesting a more stable interaction with dynein. In reciprocal experiments, c-myc-Fcho2 coimmunoprecipitated endogenous Mon1a, however, immunoprecipitation of c-myc-Fcho2 did not coprecipitate dynein (**Figure 1D**). These results suggest that Mon1a functions in two different complexes, one with dynein (1) and in an Fcho2 complex that does not interact with dynein, consistent with Mon1a functioning at multiple trafficking events.

### 3.2 FCHO2 is required for Golgi maintenance but not anterograde trafficking

We demonstrated that endocytosis was not measurably affected in Mon1a siRNA-treated cells, however movement through the secretory pathway was affected. We reported that ER to Golgi transport decreased and that Golgi stacks became less organized losing their tight ribbon structure (1). A role for FCHO2 in endocytosis has been well established but a role in the secretory pathway has not been reported. We generated an antibody to Fcho2 as described in Materials and Methods. Western analysis showed efficient RNAi-mediated knockdown of Fcho2 (**Figure 2A**). Unfortunately, our antibody did not work for immunofluorescence. In HeLa cells expressing GalNAc-T2-GFP, a resident Golgi enzyme that labels the entirety of the Golgi apparatus (21, 22), transfection with nonspecific targeting siRNAs did not affect Golgi morphology, which presented as organized perinuclear Golgi stacks with intact cisternae. In contrast, cells transfected with human FCHO2-specific siRNAs showed fragmented Golgi stacks; Golgi ribbons were disrupted into ministacks that remained centrally clustered. The fragmentation was not reminiscent of mitotic fragmentation where the Golgi becomes extensively fragmented into a “haze” of difficult to distinguish vesicles (23). We confirmed that the Golgi fragmentation was specific to silencing FCHO2 and was not due to off target effects, as HeLa cells silenced for human FCHO2 but expressing a myc-tagged siRNA-resistant Fcho2 (murine allele) did not show Golgi fragmentation (**Figure 2A**).

**Figure 2.**
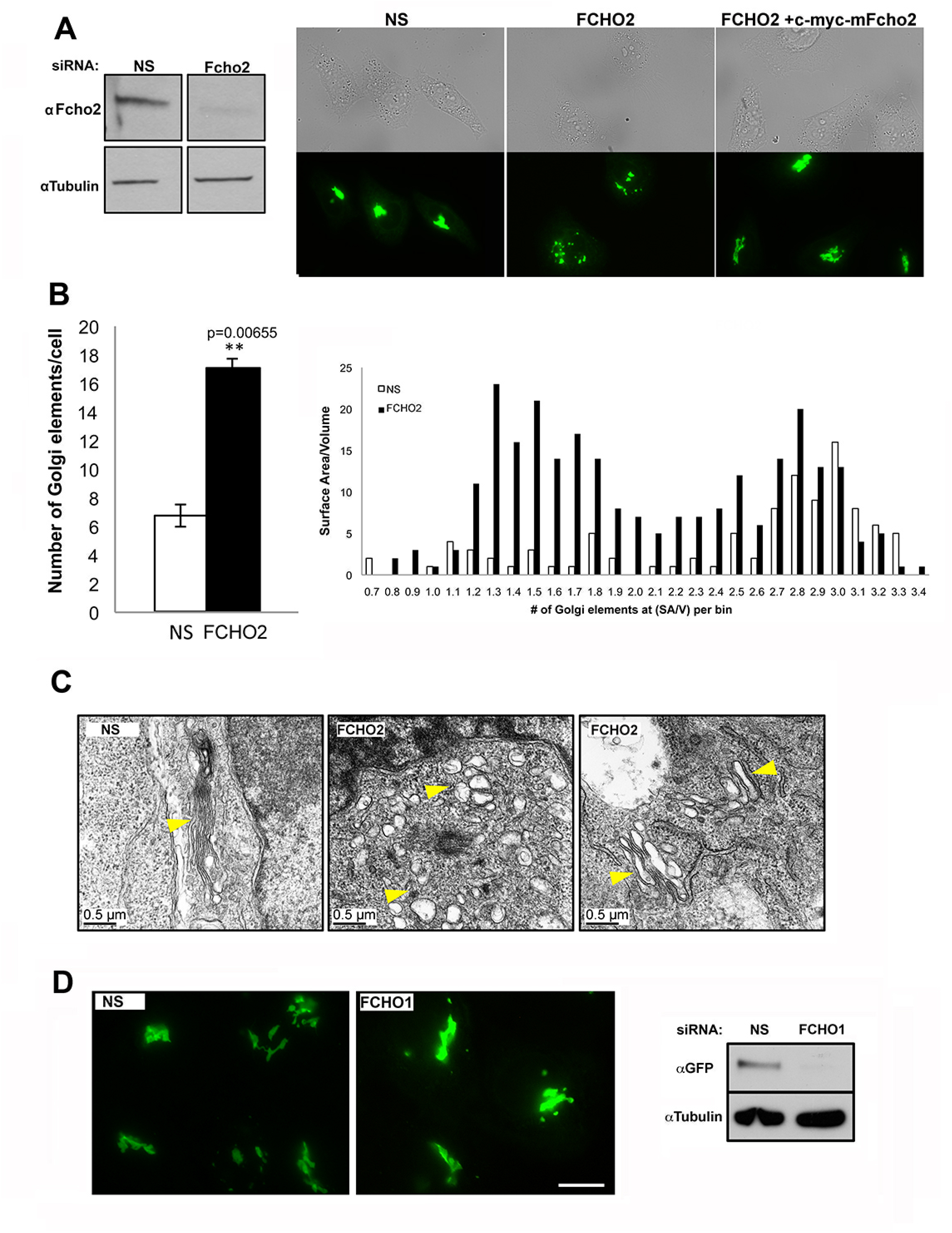
FCHO2 is required for maintenance of Golgi architecture. A. NIH3T3 cells were treated with siRNAs specific to mouse Fcho2 or nonspecific (NS) control siRNAs for 72hrs. Western analysis was done on NIH3T3 to confirm efficient reduction of Fcho2. HeLa cells expressing the Golgi protein GalNAc-T2-GFP were incubated with human FCHO2-specific siRNA for 72hrs, transfected with empty vector or pCMVTag3B c-myc-mouse Fcho2 and Golgi morphology was analyzed using live cell epifluorescence imaging of GalNAc-T2-GFP signal. B. Cells silenced as in (A) were imaged using confocal microscopy and Golgi morphology quantified using Volocity software as described previously (1). Data are expressed as Golgi elements/cell or average Golgi surface area/volume expressed as voxels. C. Cells silenced as in (A) for NS control or FCHO2-specific siRNAs were processed for electron micrograph analysis at 72hrs post RNAi-treatment. Representative images for NS (one) and FCHO2 silenced (two) cells are shown. Yellow arrows denote intact or fragmented/ministacked Golgi structures. D. HeLa cells expressing the Golgi protein GalNAc-T2-GFP as in A were incubated with NS or FCHO1 specific siRNAs for 72 hrs and imaging performed as in A. Scale bar = 5µm. HeLa cells were incubated with NS or FCHO1 specific siRNAs for 48 hrs, transfected with human GFP-tagged FCHO1 and at 72hrs post siRNA cells were lysed and GFP-FCHO1 levels determined by Western blot. A representative blot is shown (n=2).

Confocal imaging and Volocity software analysis were used to quantify the breakdown of the Golgi complex in FCHO2-depleted GalNAc-T2-GFP cells. Nonspecifically silenced cells showed tightly organized perinuclear Golgi stacks with an average of seven Golgi elements per cell (**Figure 2B**). Conversely, FCHO2-silenced cells had a fragmented Golgi with the number of Golgi elements increased to an average of 17 per cell. Measurements of FCHO2-depleted cells demonstrated a bimodal distribution of surface area to volume for Golgi fragments compared to control cells demonstrating that FCHO2-silenced cells have fewer Golgi elements with large surface area to volume ratios. This bimodal distribution may reflect cells that have yet to go through mitosis (see below). Electron micrographs of FCHO2-silenced cells also showed fragmented Golgi morphology with ministacks lacking ribbon structures (**Figure 2C**). FCHO2-silenced cells showed no changes in cell viability (data not shown).

FCHO1 is a homologue of FCHO2 and has been shown to function in endocytosis (22, 24). We examined if reductions in FCHO1 would also affect Golgi morphology. Treatment of cells with siRNA targeted against FCHO1 did not result in Golgi fragmentation (**Figure 2D**), however, we could not detect endogenous levels of FCHO1 using commercially available antibodies to confirm the efficacy of the human FCHO1 oligonucleotide pool silencing reagents (data not shown). To determine if the oligonucleotide pool against FCHO1 was effective in reducing FCHO1 levels we silenced human FCHO1 for 48 hours followed by transient expression of human GFP-FCHO1, which is predicted to be sensitive to human FCHO1 siRNA oligonucleotides. This procedure has been used previously as an indicator of siRNA efficacy when antibodies are not available (25, 26). Western blot analysis showed a marked reduction in GFP-FCHO1 (**Figure 2D**). We infer from these results that reductions in endogenous FCHO1 would be similar to reductions in overexpressed GFP-FCHO1 confirming the efficacy of the siRNA. These results suggest that FCHO1 does not function in maintaining Golgi morphology, although it is also possible that the remaining FCHO1 had residual activity.

Golgi fragmentation in FCHO2-silenced cells could result from a defect in the anterograde pathway (22, 27). To determine if FCHO2 acts in anterograde trafficking in the secretory pathway, HeLa cells expressing GalNAc-T2-GFP were transfected with FCHO2 siRNAs for 72 hours prior to treatment with the fungal metabolite Brefeldin A (BFA). BFA causes a dramatic redistribution of Golgi membrane to the ER (28); this redistribution was not affected by reduced levels of FCHO2 (**Figure 3A**). In nonspecifically silenced cells, removal of BFA allowed the Golgi to reassemble into a stacked ribbon structure within 3 hours. FCHO2-silenced cells recovered their Golgi within the same time frame as nonspecifically silenced cells, although the Golgi elements were fragmented. These data were quantified and expressed as the percent of cells showing intact or fragmented Golgi structures –BFA, ER or Golgi localization +BFA and ER, intact or fragmented Golgi for the three hour BFA recovery (**Figure 3B**). We conclude that ER to Golgi transport is not altered in FCHO2-silenced cells. This is different from Mon1a silencing and suggests that the FCHO2-Mon1a interaction functions at a different site.

**Figure 3.**
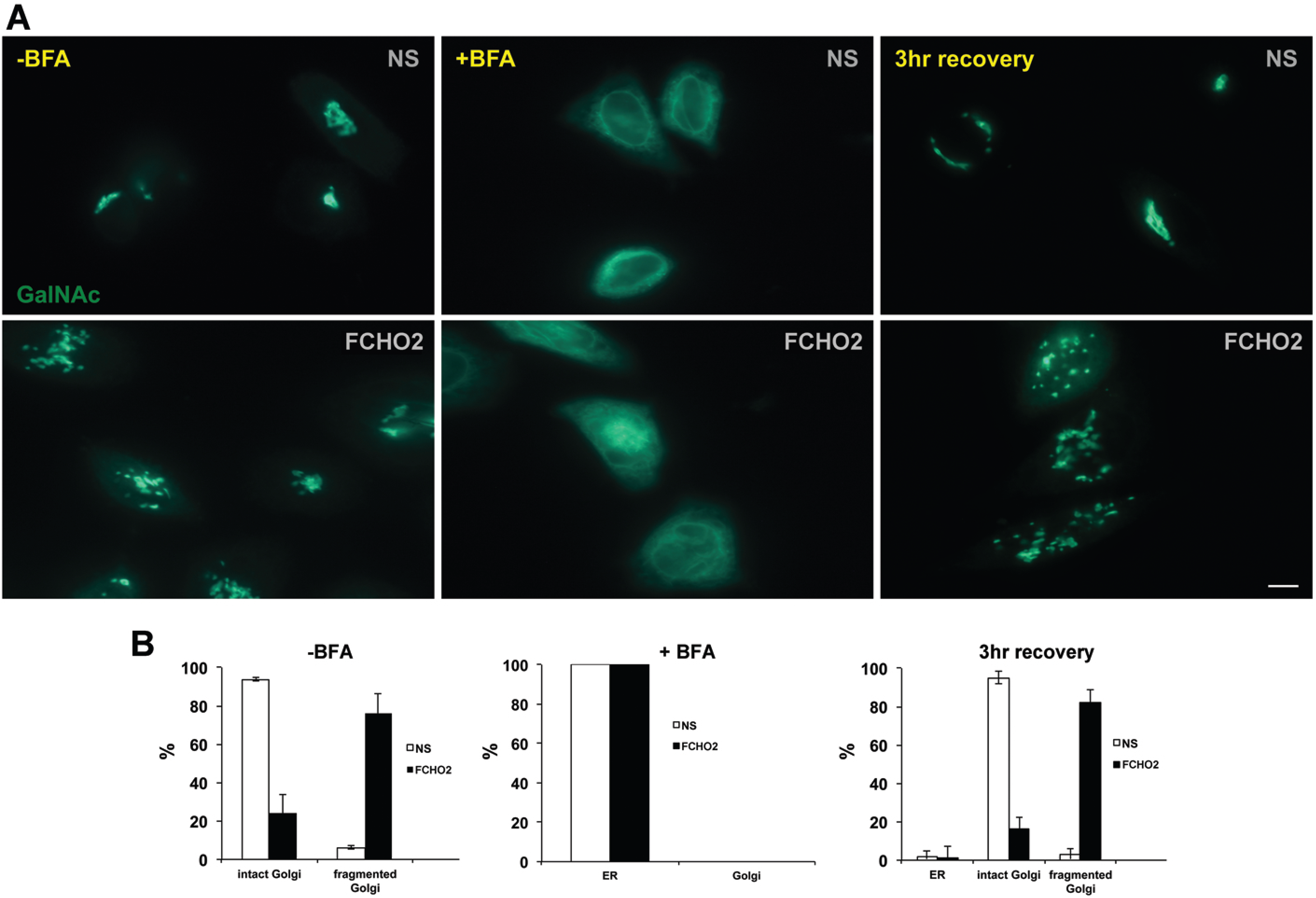
FCHO2 is not required for ER-Golgi transport. A. HeLa cells stably expressing GalNAc-T2-GFP were treated with siRNAs as described in Figure 2. Silenced cells were then incubated with BFA (5 μg/mL) for 30 min at 37°C before stringent washing and incubation for three hrs in the absence of BFA. Movement from ER to Golgi was assessed by live cell epifluorescence microscopy. B. Images from two separate experiments (>50 cells/treatment) were quantified for intact or fragmented Golgi –BFA (left panel), ER or Golgi localization of GalNAc-T2-GFP after BFA (middle panel) and ER, intact or fragmented Golgi after three hr recovery from BFA treatment (right) panel. All numbers were normalized to (-) BFA where 100% of the cells showed ER localization. Error bars represent the SD of n=2.

### 3.3 siRNA targeting Mon1a (5’-UTR) results in Golgi fragmentation

We reported that silencing of Mon1a affected ER to Golgi traffic yet only had a mild effect on Golgi morphology, the Golgi appeared less organized but the cisternae were intact (1). The finding that Mon1a interacts with FCHO2 and that reductions in FCHO2 lead to Golgi fragmentation led us to reexamine the effect of Mon1a silencing on Golgi morphology. We considered the possibility that the residual Mon1a protein remaining after RNAi treatment was sufficient for Golgi complex maintenance. Previously, we utilized a siRNA pool strategy to reduce Mon1a levels, using oligonucleotides targeted to sequences in the open reading frame (ORF). To determine if we could reduce Mon1a levels more efficiently, RNAi oligos were designed to target either the ORF or the 5’-untranslated region (5’-UTR) of Mon1a. HeLa cells expressing GalNAc-T2-GFP were treated with each siRNA for 72 hours and Golgi morphology was assayed by epifluorescence microscopy. RNAi oligos specific to a sequence in the ORF of Mon1a led to the same Golgi phenotype as previously published, the Golgi was disorganized yet remained intact (1). Surprisingly, cells treated with oligos targeting the 5’-UTR of Mon1a led to fragmented Golgi stacks that were similar to that seen in FCHO2-silenced cells as assayed by either fluorescence (**Figure 4A**) or electron microscopy (**Figure 4B**). Mon1a silenced cells did not show reductions in cell viability (data not shown). Quantification of Mon1a protein levels showed that the 5’-UTR targeted oligos reduced Mon1a protein levels at 48 hours more than the ORF targeted oligos. At 72 hours the decreases in protein levels were the same for both the 5’-UTR and the ORF targeted oligos (**Figure 4C**). We confirmed that ER to Golgi trafficking in Mon1a (5’-UTR) silenced cells was impaired, similar to our previous findings (1). Importantly, the defect in ER to Golgi traffic, as well as Golgi disruption, was suppressed when Mon1a (5’-UTR) silenced cells were transfected with a plasmid expressing a silencing-resistant FLAG-Mon1a (**Figure 4D**). These results show that the fragmented Golgi seen in Mon1a (5’-UTR) silenced cells is not due to an off target effect of the 5’-UTR silencing oligo. These results also confirm that Mon1a is acting at the Golgi apparatus to maintain Golgi ribbon morphology.

**Figure 4.**
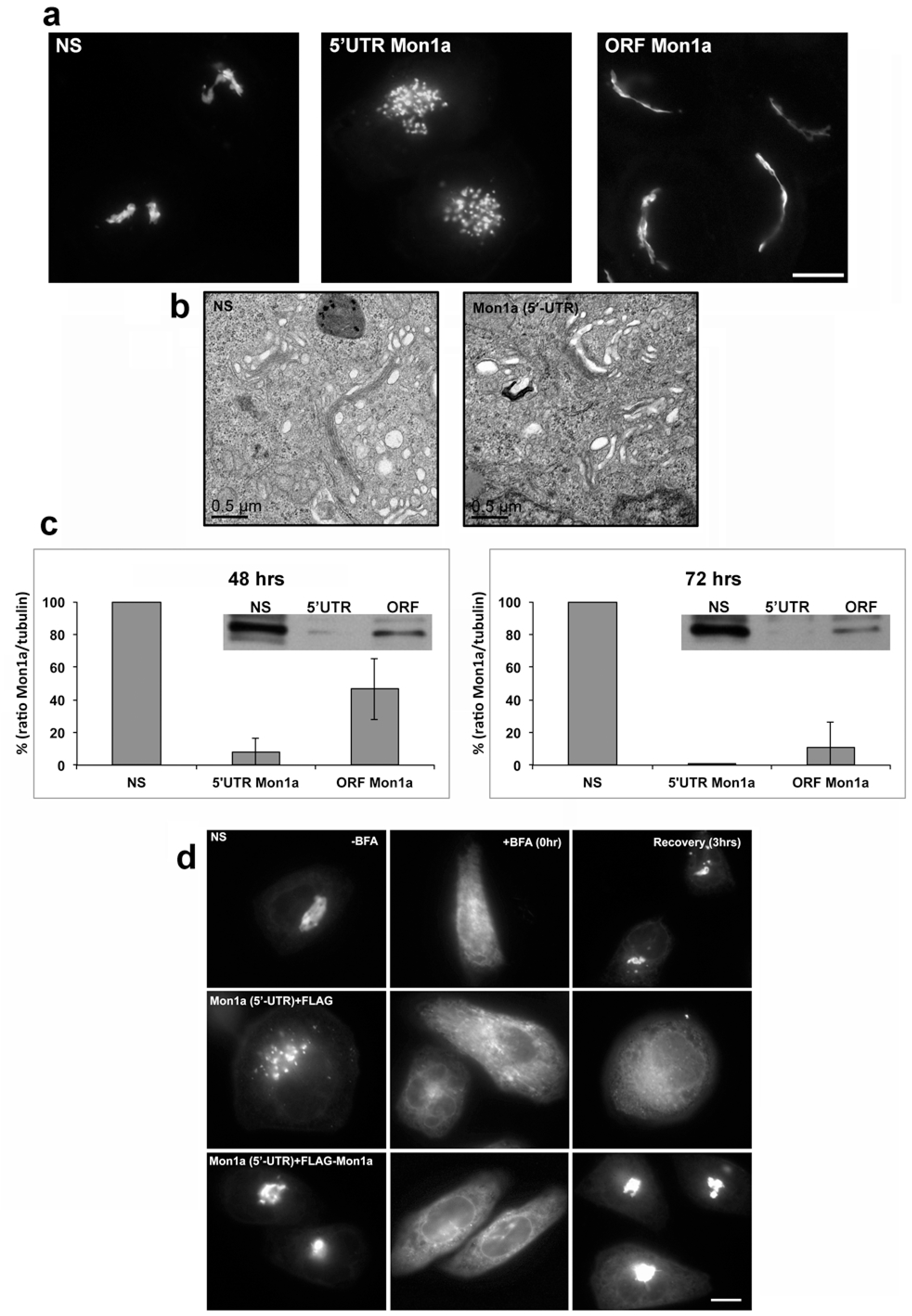
siRNA targeting 5’-UTR of Mon1a fragments the Golgi apparatus. A. GalNAc-T2-GFP expressing HeLa cells were transfected with NS, 5”UTR Mon1a or ORF Mon1a and Golgi morphology examined using epifluorescence microscopy. Scale bar = 5µm. B. siRNA transfected cells were fixed at 72 hours of silencing and ultrastructural analysis was performed. Representative electron micrographs are shown. C. Cells silenced as in A for 48 or 72 hrs were lysed and Mon1a and tubulin levels determined by Western blot. Blots were quantified using BioRad gel quantification software and the amount of Mon1a/tubulin determined. Error bars represent the standard deviation of the mean. Included in the graph are representative westerns of Mon1a silencing. D. Mon1a-silenced cells were transfected with FLAG-Mon1a or an empty vector control and Golgi morphology and ER-Golgi trafficking using a BFA recovery assessed by epifluorescence microscopy.

### 3.4 Co-depletion of Rab6 and FCHO2 or Mon1a suppresses silencing-induced Golgi fragmentation

Reductions in the levels of a number of different proteins can affect Golgi architecture and function. Loss of these proteins disrupts Golgi structure by affecting vesicle fission from a donor compartment or by affecting fusion of vesicles with recipient compartments. Two examples are the Golgi GTPase Rab6 and the ER tether ZW10 /RINT-1, which are required for vesicle fission and fusion respectively to maintain the fidelity of membrane trafficking and Golgi morphology (22). RNAi-dependent knockdown of the retrograde tether ZW10/RINT-1 resulted in a central clustering of Golgi fragments (22). Rab6 is an evolutionarily conserved GTPase known to facilitate trafficking from the Golgi to the cell surface (anterograde), intra-Golgi transport mediated by the retrograde tether complex COG and trafficking from the Golgi to the ER (retrograde) through a physical interaction with dynein (22, 29, 30). Rab6 functions are epistatic to those of ZW10; co-silencing results in a Rab6 phenotype with intact Golgi architecture (22). To determine if Rab6 was epistatic to the Golgi fragmentation phenotypes associated with siRNA knockdown of FCHO2 or Mon1a, GalNAc-T2-GFP expressing HeLa cells were treated with siRNA oligonucleotides specific to Mon1a (5’-UTR), FCHO2 or Rab6, alone or in combination. Control and Rab6 depleted cells showed intact Golgi cisternae while Mon1a (5’-UTR) and FCHO2 silenced cells possessed fragmented Golgi elements as previously demonstrated (**Figure 5A**). Co-silencing of Rab6 and FCHO2 or Mon1a (5’-UTR) preserved the morphology of the Golgi stacks resulting in a Rab6 silencing phenotype as opposed to a fragmented morphology. The percentage of cells showing Golgi fragmentation was quantified (**Figure 5B**). That depletion of Rab6a suppresses the effects of Mon1a and FCHO2 silencing demonstrates that Rab6 is epistatic to Mon1a or FCHO2 in Golgi morphology maintenance.

**Figure 5.**
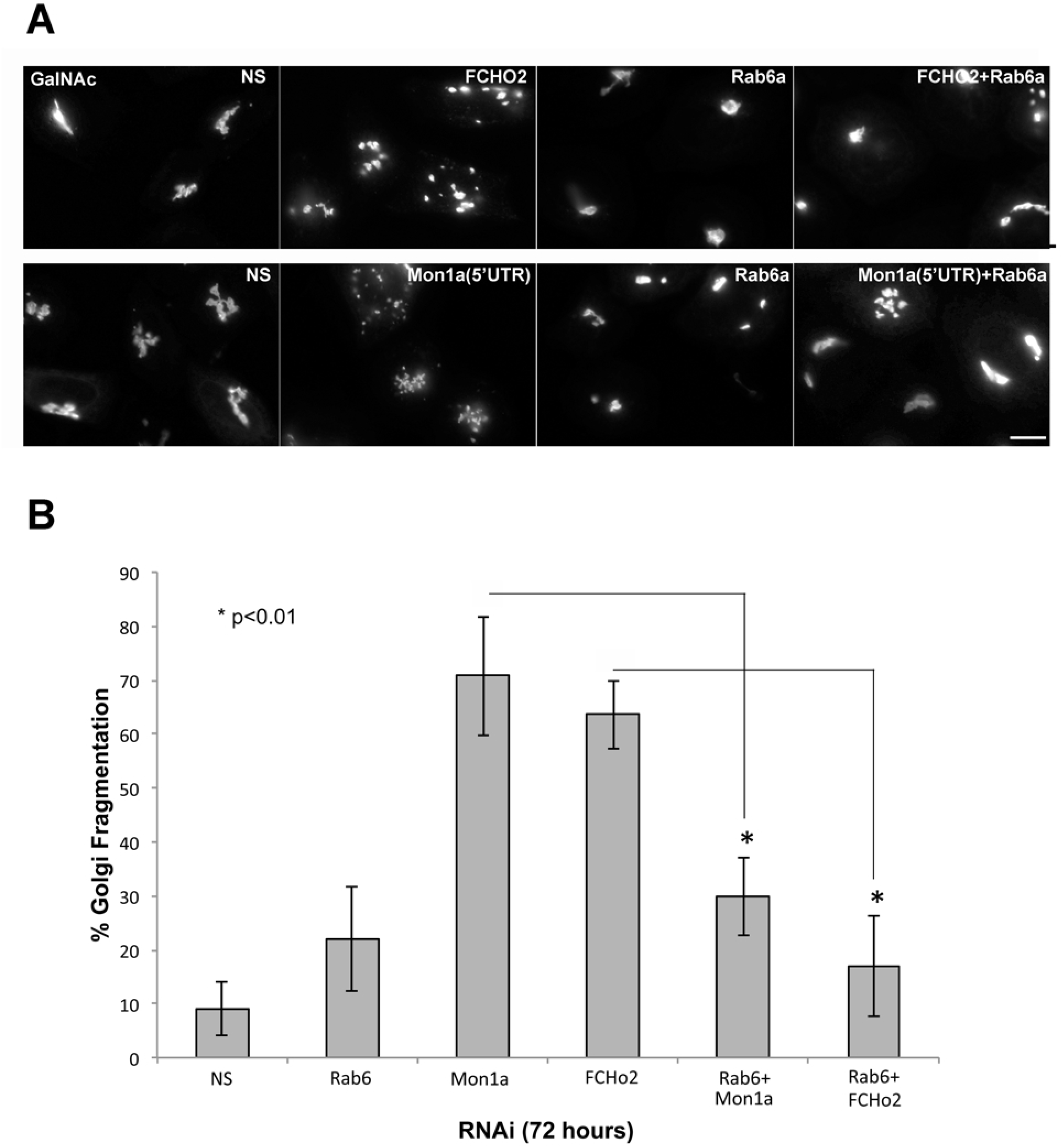
Rab6 is epistatic to Mon1a and FCHO2 silencing-induced Golgi fragmentation. A. HeLa cells expressing GalNAc-T2-GFP were silenced with Mon1a, FCHO2 or Rab6 alone or in combination for 72 hours. Epistasis was established by scoring Golgi morphology using live cell epifluorescence microscopy. B. Quantification of four independent epistasis experiments showing percent Golgi fragmentation with p values less than 0.01 designated (*).

### 3.5 FCHO2 depletion does not affect trafficking through the secretory pathway

Studies have shown that Golgi fragmentation does not necessarily impair secretion (19, 31). Our data show that ER to Golgi trafficking in FCHO2-silenced cells is unaffected. To determine if reductions in FCHO2 affect Golgi to PM trafficking we took advantage of a temperature-sensitive allele of VSVGtsGFP that concentrates and restricts its localization to the ER at the restrictive temperature (39°C). Shifting cells to the permissive temperature (32°C) allows cells to traffic VSVGtsGFP from ER to Golgi and plasma membrane. An intrinsic trait of all mammalian cells is that membrane traffic is temperature-sensitive. Incubation of cells at 20°C allows trafficking from the ER to the Golgi but delays trafficking to the plasma membrane. To examine Golgi-PM transport in FCHO2-silenced cells VSVGtsGFP was allowed to accumulate at the Golgi at 20°C before shifting to 32°C. The movement of VSVGtsGFP out of the Golgi to the cell surface was unimpaired in FCHO2-depleted cells (**Figure 6A**). These data demonstrate that the fragmentation of the Golgi seen in FCHO2-silenced cells did not impair trafficking through the secretory pathway.

**Figure 6.**
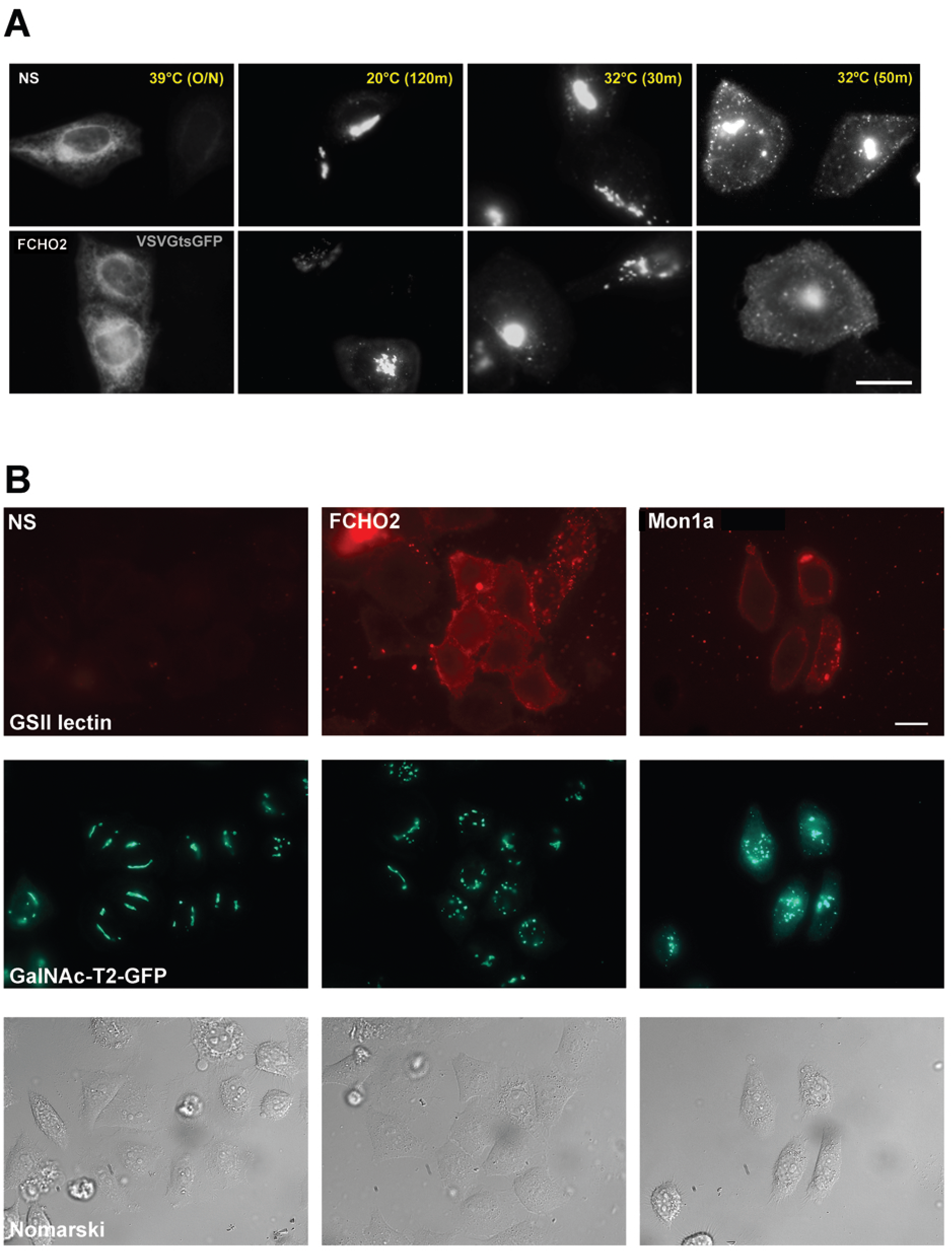
FCHO2-silenced cells show normal kinetics of Golgi to cell surface trafficking but increased immature glycosylation structures. A. HeLa cells transfected with nonspecific or FCHO2-specific siRNA for 48 hours and then cells were transfected with VSVGtsGFP and grown overnight at 39.5°C. Cells were shifted to 20°C for two hours allow VSVGtsGFP capture in the Golgi apparatus. Cells were then shifted to the permissive temperature (32°C) and images captured at 30 and 50 minutes. Representative images are shown from >50 cells per treatment. N=3. B. GalNAc-T2-GFP cells were incubated with NS, FCHo2 or Mon1a siRNA for 72 hrs, cells placed at 4°C and washed with Hanks MEM + 10% FBS and incubated at 4°C with the cell impermeable Alexa594-conjugate GS-II lectin for 30 mins. Cells were extensively washed and epifluorescence images captured. Representative images are shown from >50 cells. N=2

Carbohydrate processing requires the sequential activity of a number of enzymes that span the entirety of the Golgi stack. Glycosylation of proteins can be impaired when properly organized Golgi architecture is disrupted(19, 32). N-acetyl-D-glucosamine is a carbohydrate structure rarely seen at the plasma membrane because it is processed and modified by enzymes in the organized Golgi stack. When Golgi architecture is disrupted N-acetyl-D-glucosamine can be inappropriately found at the cell surface(19). We examined carbohydrate processing by staining live cells with Alexa594-conjugated lectin (GS-II). This lectin binds to terminal N-acetyl-D-glucosamine residues. In mature glycoproteins these residues are not found as terminal carbohydrates and cells show little staining with Alexa594-conjugated lectin (GS-II). In cells with disruptions in terminal carbohydrate modifications there is an increase in lectin binding. FCHO2-silenced cells had increased GS-II binding at the cell surface while control cells had almost no GS-II fluorescence (**Figure 6B**). Previously, we determined that ER to Golgi trafficking and Golgi to plasma membrane trafficking of the temperature sensitive allele of VSVG-GFP was reduced when cells were silenced for Mon1a (1). We did not examine if there were changes in carbohydrate modifications in that study. Similar to FCHO2 silenced cells, Mon1a-silenced cells showed increased GS-II staining (**Figure 6B**). These results show that the fragmentation of the Golgi apparatus in FCHO2- or Mon1a-silenced cells gives rise to impaired carbohydrate processing and suggests an important role for FCHO2 and Mon1a in maintaining Golgi architecture.

### 3.6 Reductions in FCHO2 or Mon1a affect lateral Golgi transfer

The above data shows that disrupted Golgi in FCHO2-silenced cells can still vectorally transfer molecules through the secretory pathway; yet, Golgi architecture is altered with the notable absence of Golgi ribbons that connect Golgi stacks. A decrease in ribbon formation would affect lateral transfer between Golgi stacks. We utilized fluorescence recovery after photobleaching (FRAP) to determine if there is communication between Golgi elements in FCHO2 or Mon1a-silenced cells (19, 31, 33, 34). FCHO2-or Mon1a-silenced GalNAc-T2-GFP expressing cells were imaged with epifluorescence confocal microscopy. A region of the Golgi structure was bleached and FRAP monitored. FRAP occurred in a matter of seconds in nonspecifically-silenced cells (**Figure 7A,B**). Conversely, recovery was greatly reduced in Mon1a and FCHO2-silenced cells. These data suggest that Mon1a and FCHO2 are both required for membrane ribbon formation and intra-Golgi communication.

**Figure 7.**
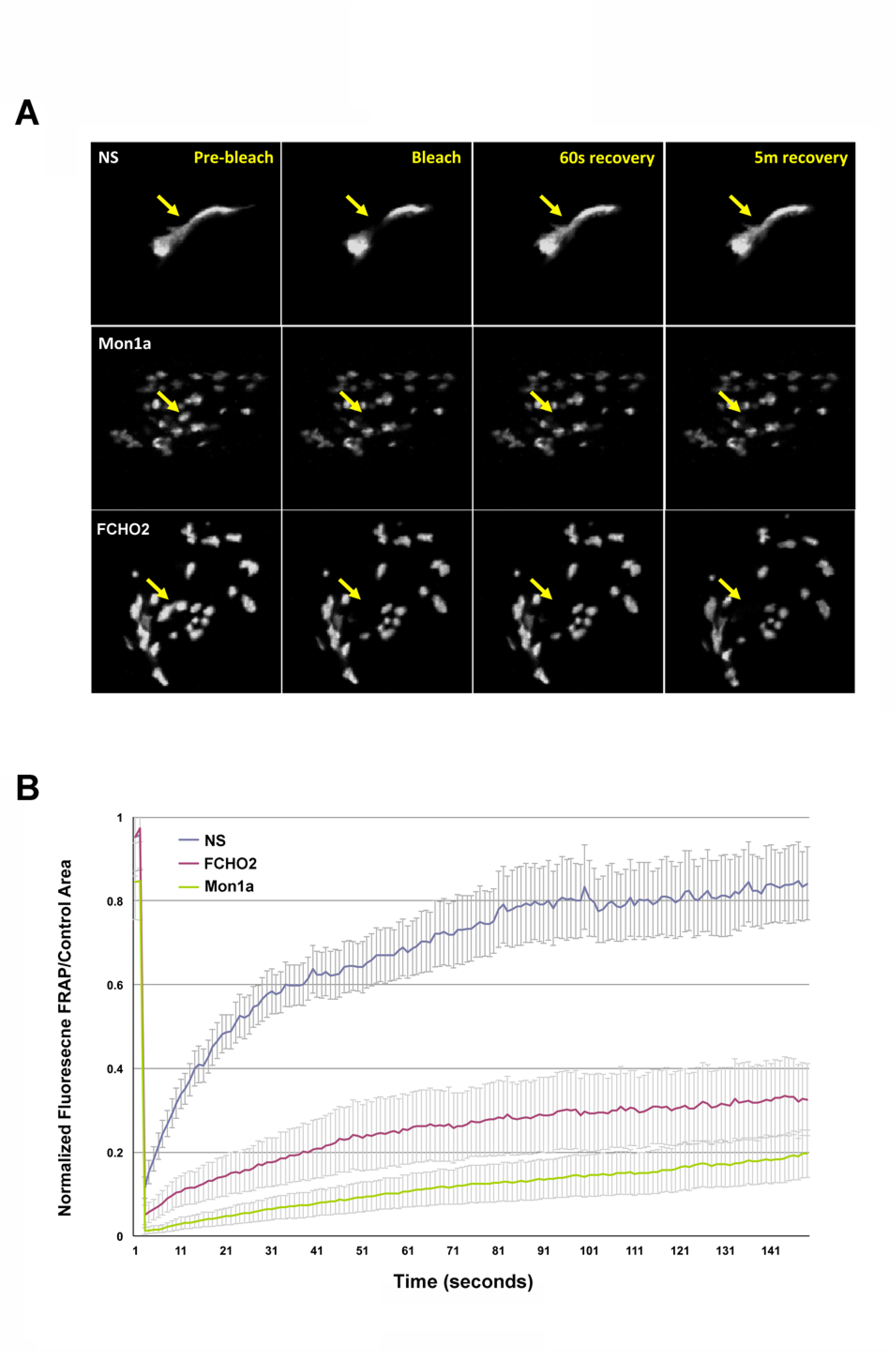
Golgi FRAP is reduced in Mon1a and FCHO2-silenced cells. A. Cells treated with nonspecific, FCHO2 or Mon1a oligos for 72 hours were visualized by live cell microscopy. Golgi elements (GalNAc-T2-GFP-positive) were photobleached and recovery assessed at indicated times by time-lapse microscopy. An example set of images is shown. B. Data from three separate experiments as in A were analyzed using eight-ten cells each, the mean determined and the data presented in graph form +/- SEM. Fluorescence recovery was determined by dividing the GFP fluorescence in the bleached spot by that in a nearby unbleached region as described (19).

### 3.7 Golgi fragmentation in FCHO2-depleted cells is cell cycle-dependent

Golgi material is partitioned between daughter cells during mitosis, which is facilitated through the systematic breakdown of Golgi ribbons to individual stacks and ultimately into vesicles. Following mitosis, Golgi vesicles coalesce and the organelle is reassembled. Golgi ministacks are reformed and then the ministacks are connected by membrane ribbons (35, 36). There are two likely mechanisms that might explain the disruption of Golgi ribbons in FCHO2-silenced cells. First, ministack fragments in FCHO2-silenced cells may result from ribbon breakdown, as loss of FCHO2 destabilizes Golgi ribbons. If FCHO2 is required for ribbon stability, then reducing FCHO2 levels should be sufficient to induce Golgi fragmentation. Alternatively, FCHO2 may act after mitosis when Golgi reformation occurs by tubulating Golgi membranes into Golgi ribbons permitting those ribbons to link individual stacks. RNAi and cell cycle arrest were utilized to address if FCHO2 or Mon1a activity was required in Golgi ribbon stability versus ribbon formation. If loss of FCHO2 is required for ribbon maintenance, then the loss of Golgi ribbons should be independent of mitosis. In contrast, if Golgi ribbon formation requires FCHO2, then inhibition of mitosis should prevent the siRNA-dependent ribbon disruption. To test this hypothesis, Mon1a, FCHO2 and dynein heavy chain1 were knocked down using siRNA in cell cycle-arrested GalNAc-T2-GFP-expressing HeLa cells. Incubation in thymidine blocks cells at the G1/S phase arresting the cell cycle before mitosis preventing Golgi disassembly. Cells incubated with nonspecific siRNA possessed tightly organized Golgi stacks in the presence or absence of thymidine (**Figure 8A,B)**. Silencing of dynein heavy chain1 resulted in Golgi disruption in both thymidine treated and untreated cells. This result indicates that thymidine treatment does not prevent Golgi fragmentation seen in dynein-silenced cells. Golgi fragmentation in Mon1a-silenced cells also was not affected by thymidine treatment. In contrast, thymidine treatment affected Golgi fragmentation in FCHO2-silenced cells. There was greater than 50% reduction in fragmented Golgi in cells blocked at G1/S. These results suggest that Mon1a and dynein may affect multiple steps required for Golgi architecture, whereas FCHO2 is involved in ribbon formation during Golgi reassembly after mitosis.

**Figure 8.**
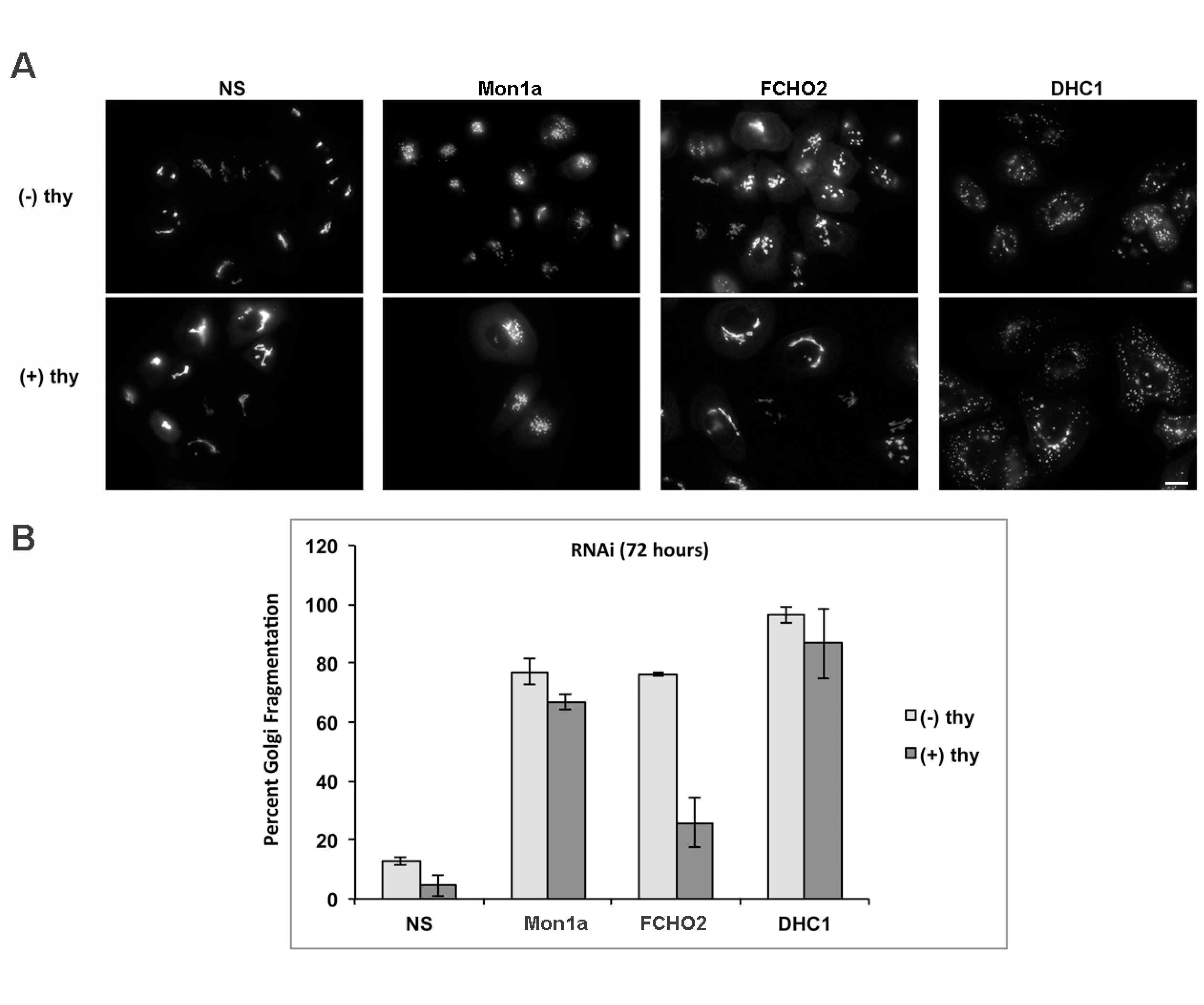
Golgi fragmentation in FCHO2-depleted cells is cell cycle-dependent. A. GalNAc-T2-GFP expressing cells were transfected with siRNA nonspecific, Mon1a, FCHO2 or DHC1 for 24 hours before cell cycle arrest was achieved by 2 mM thymidine treatment for an additional 48 hrs. Golgi morphology was scored using epifluorescence microscopy with representative images shown. B. Cells showing Golgi fragmentation were quantified from two independent experiments with over 100 cells per condition score and the results graphed +/- standard deviation.

### 3.8 FCHO2 is localized to the Golgi apparatus in a Mon1a-independent fashion

To determine if FCHO2 is localized to the Golgi apparatus and if Mon1a is required for that localization we generated a mCherry-FCHO2 plasmid. We confirmed that this plasmid complements the siRNA dependent FCHO2 depletion of fragmented Golgi seen in GalNAc-T2-GFP-expressing HeLa cells (data not shown). We used time-lapse spinning disk confocal microscopy to image Golgi morphology and FCHO2 localization in nonspecifically silenced and Mon1a-silenced cells. mCherry-FCHO2 was found colocalized with intact Golgi ribbons in nonspecifically silenced cells (**Figure 9A**) as well as with Golgi fragments seen in Mon1a-silenced cells (**Figure 9B**). Time-lapse microscopy showed that the presence of mCherry-FCHO2 on GalNAc-T2-GFP Golgi structures was transient with colocalization sites changing over time (**Figure 9 arrows and Supplemental Movies 1,2**). These data demonstrate that FCHO2 frequently interacts with the Golgi apparatus and that this interaction is not dependent upon the presence of Mon1a. This also suggests that Mon1a acts in Golgi ribbon formation after the action of FCHO2.

**Figure 9.**
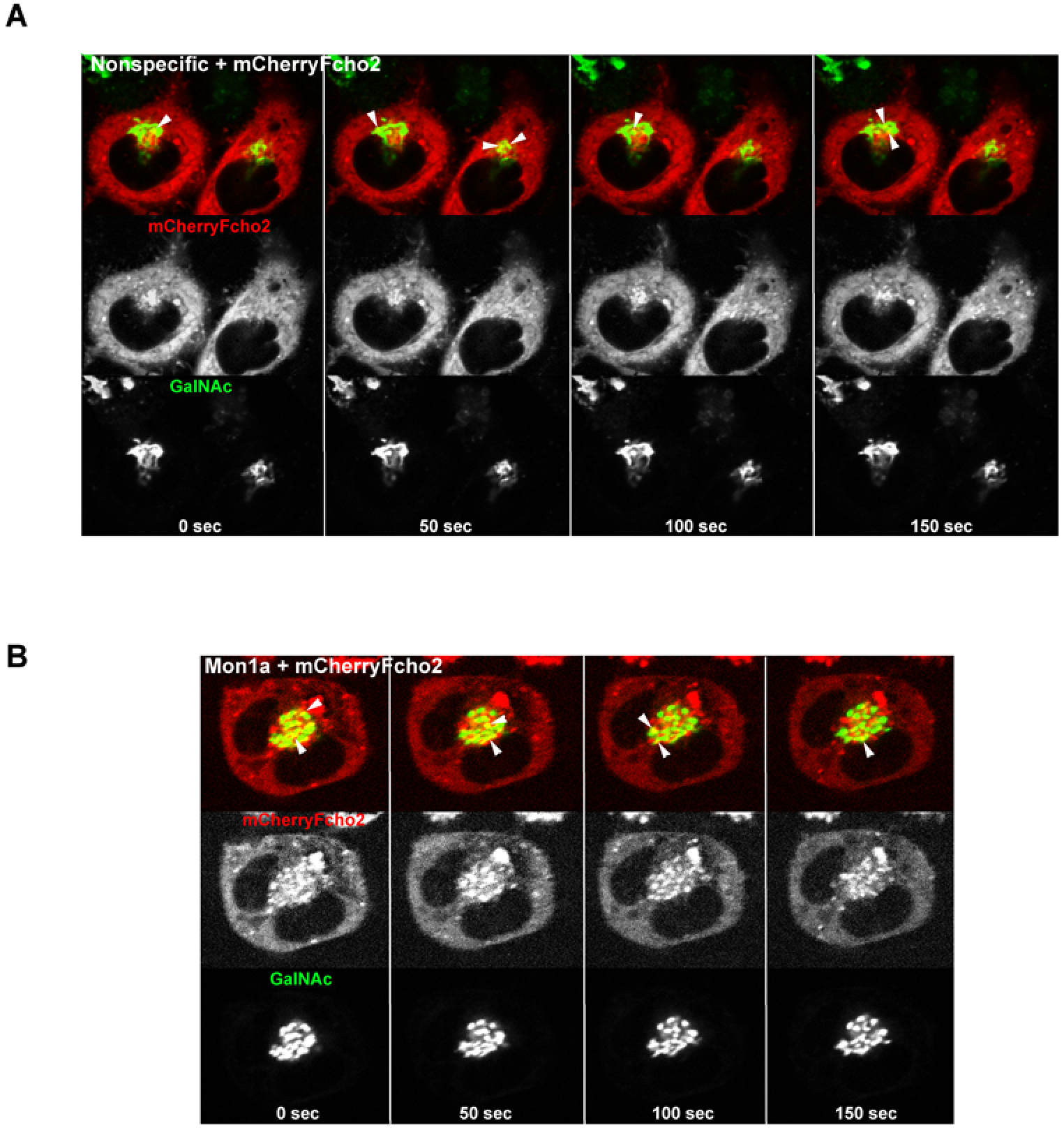
mCherry-FCHO2 localizes to the Golgi apparatus in the absence of Mon1a. HeLa cells expressing GalNAc-T2-GFP were silenced with nonspecific (A) or Mon1a (B) oligonucleotides for 72 hours. Cells were then transfected with mCherry-FCHO2 and live cell imaging captured using a Spinning Disk Confocal Microscope. Images were captured at a frame per second with a Z series captured every 10 seconds. To examine colocalization with GalNAc-T2-GFP Golgi, the Golgi focal plane was used to generate the still frames. Representative time-frame images are shown every 50 seconds with the merge (top panel), mCherry-FCHO2 (middle panel) and GalNAc-T2-GFP (Bottom panel). Arrows denote colocalization sites and changes in colocalization sites over time.

## 4. Discussion

Our previous studies showed that Mon1a played a role in vesicle formation at the ER and interacted with the microtubule motor protein dynein involved in vesicle trafficking at the Golgi (1). Here we determined that Mon1a associates with a higher molecular weight complex that contains more than one copy of Mon1a. Using two-hybrid and coimmunoprecipitation analysis we identified FCHO2 as a Mon1a-interacting protein. FCHO2 is a member of a large family of proteins known as F-BAR domain-containing proteins, many of which have been demonstrated to function in endocytosis (20, 37–41). This role in endocytosis requires the F-BAR domain of FCHO2 and its membrane bending capacity, suggesting that FCHO2 deforms and tubulates membranes as shown *in vitro* (42). We determined that reduced levels of FCHO2 resulted in fragmentation of the Golgi apparatus but ER to Golgi trafficking was unaffected. Previous studies suggested that the F-BAR protein CIP4 is associated with the Golgi apparatus and that reductions in AKAP350, an A kinase anchoring protein that binds CIP4, lead to changes in Golgi structure (43). Those studies, however, did not look at reductions in CIP4 and its effect on the Golgi morphology. Microscopic analysis revealed that FCHO2 silencing resulted in Golgi ministacks, a morphology often found after mitosis as Golgi vesicles re-fuse and reconnect ministacks (44). The finding that inhibition of mitosis reduced FCHO2-induced Golgi fragmentation suggests that FCHO2 is required for the reformation of the Golgi ribbon after mitosis, a process that might require significant membrane tubulation. This role is consistent with the identified role of FCHO2 as a membrane deforming and tubulating protein (43, 45). This is the first report that reductions in the F-BAR protein FCHO2 affect Golgi morphology.

Our previous study suggested that reductions in Mon1a affected vesicle formation at the ER, that Mon1a interacted with dynein and that trafficking through the secretory pathway was delayed in Mon1a-silenced cells, although those studies did not show evidence of Golgi fragmentation (1). In our current study we determined that more efficient silencing of Mon1a resulted in Golgi fragmentation. Silencing using oligonucleotides directed at the 5’-UTR of Mon1a was more efficient in reducing Mon1a levels compared to oligonucleotides directed at the Mon1a ORF. Our previous studies and current results also demonstrate that Mon1a acts in two different complexes: 1) a complex with the microtubule motor protein dynein to effect trafficking through the secretory pathway (1) and 2) a complex with FCHO2 involved in maintaining Golgi architecture. We note that the effects of different levels of Mon1a silencing are consistent with the strength of the biochemical interaction observed. That is, less efficient reductions in Mon1a revealed a role for Mon1a in ER-Golgi trafficking, a dynein-requiring process that showed a strong biochemical interaction, but perhaps did not disrupt the Mon1-FCHO2 interaction, thus leaving the Golgi intact.

We determined that FCHO2’s role in Golgi ribbon formation is cell cycle-dependent but we did not observe any cell cycle dependence for Mon1a or dynein siRNA-dependent Golgi fragmentation. That Mon1a acts in two different complexes in the mammalian secretory pathway suggests that interpreting the resulting phenotypes associate with its loss may be compounded by affecting two different trafficking events.

Golgi fragmentation has been reported when vesicle docking proteins such as p115 have been depleted (46–48), Rab proteins have been mutated (49) or if the ADP-ribosylation factor guanine nucleotide exchange factor (ARF-GEF) ARNO3 has been overexpressed (50). p115 is essential for biogenesis of the Golgi apparatus and is an example of a protein that exhibits multiple interactions with proteins important in ER and Golgi vesicle trafficking including Rab1 at the ER and golgins at the Golgi. p115 is suggested to enhance vesicle tether complex formation and maintain Golgi morphology (51, 52). We have shown that Mon1a is important in vesicle formation at the ER, in maintenance of Golgi architecture and trafficking through the secretory pathway similar to p115. Several Rabs have been shown to function in ER and Golgi trafficking (53, 54). Rab6 has been characterized as an important regulator of membrane trafficking in the secretory pathway (22, 29, 55–58). Recently, a homologue of Rab6, Rab41, was identified to be involved in maintaining Golgi morphology and silencing Rab41 resulted in Golgi fragmentation (59). Rab41 function is epistatic to Rab6 function as silencing both Rab41 and Rab6 resulted in a Rab41 silencing phenotype of fragmented Golgi. Using epistasis analysis we confirmed that reductions in Rab6 did not affect Golgi morphology, however, reductions in Rab6 prevented the fragmentation of the Golgi apparatus seen in both Mon1a- and FCHO2-silenced cells. These results confirm that the action of Rab6, Golgi fragmentation, occurs prior to the action of Mon1a or FCHO2 in Golgi morphology maintenance. The yeast and plant Mon1 homologue, of which there is only one compared to two Mon1 proteins in mammals, complexes with Ccz1 to act as a GEF for the late endosomal Rab Ypt7 (6, 60). To date we have not identified by mass spectrometry or two-hybrid analysis, a Rab or ARF protein interacting with mammalian Mon1a. Identifying other interacting proteins in the Mon1a complex and Mon1a domains necessary for dynein or FCHO2 interactions will elaborate on mechanistic details of Mon1a function.

We identified FCHO2 as a Mon1a-interacting protein and that Golgi fragmentation associated with reductions in FCHO2 is cell cycle-dependent. The importance of fragmenting the Golgi apparatus for inheritance into daughter cells has been established (44, 61, 62). Linking the Golgi apparatus into a ribbon structure has been suggested to be important to accommodate complex protein glycosylation and alterations in this process can affect development and result in disease states (63–67). Our data confirms this point, we observed increased immature glycosylation moieties in FCHO2 and Mon1a silenced cells. Several proteins have been identified to be important in linking Golgi ministacks. For example, the Golgi reassembly stacking proteins (GRASPs) GRASP55 and GRASP65 have been described to regulate ribbon formation and participate in controlling cell cycle progression (19, 46, 68–70). These proteins have been shown to interact with Golgi tethering molecules including p115 and assist in Golgi stacking.

Based upon this current study, we have developed a model where Mon1a transiently interacts with FCHO2 at the Golgi in tubulation and vesicle formation to reconnect Golgi ministacks after mitosis (**Figure 10**). We have determined that FCHO2 is recruited to Golgi membranes in the presence or absence of Mon1a and appears to frequently traffic to and from the Golgi. Using epitope-tagging, we were unable to detect Mon1a and FCHO2 colocalization at the Golgi apparatus (data not shown). This suggests that FCHO2 may act first in Golgi ribbon formation after mitosis and Mon1a acts in a later step, perhaps in vesicle formation similar to its role in ER to Golgi trafficking (1). We do not know how FCHO2 is recruited to Golgi membranes. FCHO2 has been characterized to bind phosphoinositides such as phosphatidyl serine and phosphatidylinositol 4,5-bisphosphate (20, 38, 42, 43). Similarly, the Mon1-Ccz1 complex in *S. cerevisiae* and *C. elegans* is recruited to membranes via phosphoinositides (3, 71, 72). These lipids or other adaptor molecules, such as Eps15, similar to recruitment to the plasma membrane or endosomes, may mediate recruitment of FCHO2 and Mon1a to the Golgi.

**Figure 10.**
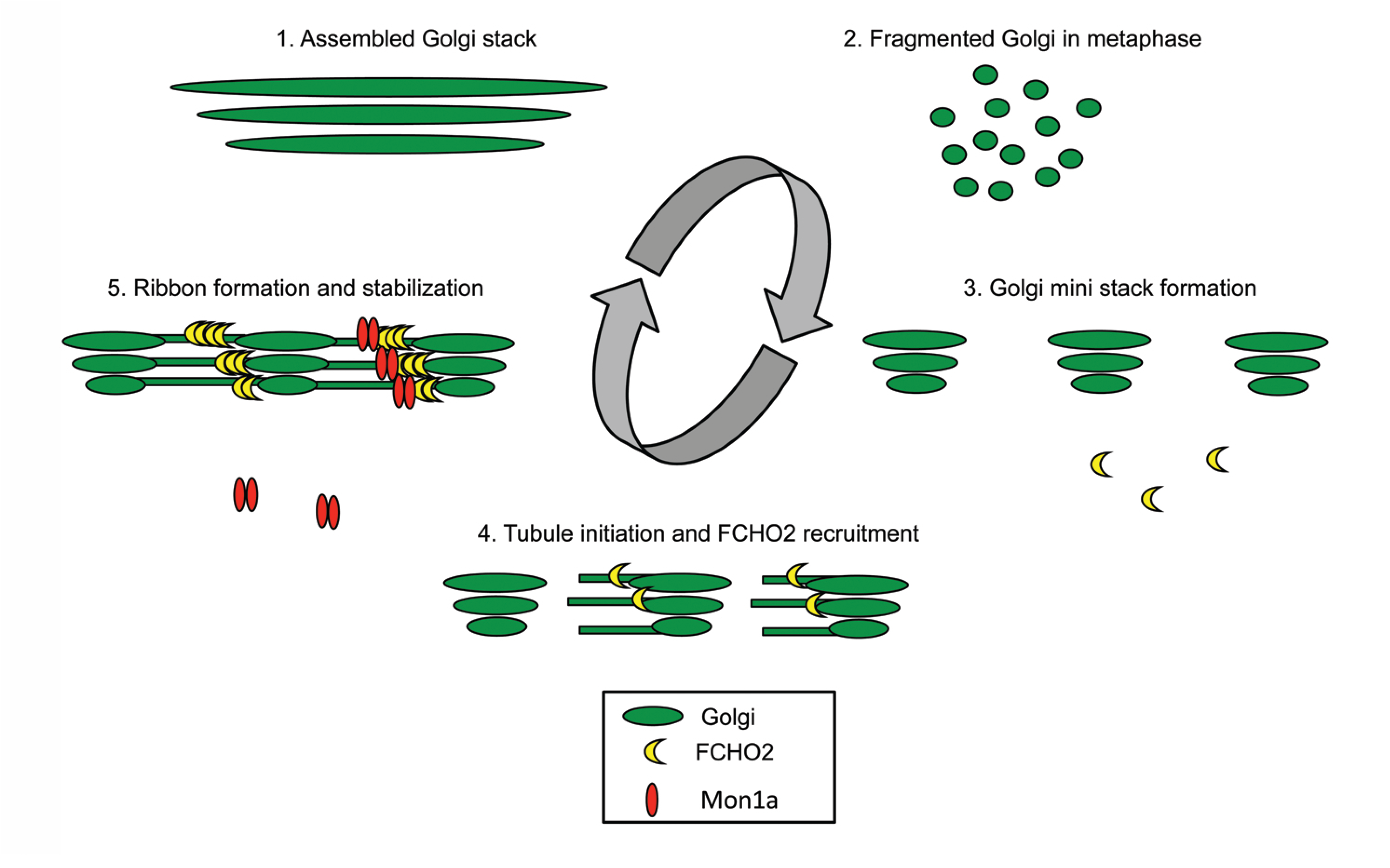
Model of Mon1a and FCHO2 function in Golgi ribbon/stack formation. 1. The Golgi forms a stack of sequential ribbon compartments that communicate through vesicle fusion and fission events. 2. During mitosis the Golgi is fragmented into ministacks and a vesicle “haze”. 3. The Golgi reforms after mitosis by first establishing ministacks. 4. FCHO2 is recruited to Golgi membranes to extend tubules to reestablish Golgi ribbons. 5. Mon1a assist in Golgi membrane tubulation and Golgi ribbon stabilization.

Our confirmation of the two-hybrid analysis, that Mon1a and FCHO2 interact in a complex, and that significant reductions in either of these proteins affect the morphology of the Golgi apparatus is a new finding. Our results that fragmentation of the Golgi in FCHO2-silenced cells is cell cycle dependent but the Mon1a-silencing Golgi fragmentation is not, is confusing as to the function of the Mon1a-FCHO2 interaction. Further, we still don’t know the stoichiometry of the FCHO2-Mon1a interaction. One caveat that must be considered in all of these results is that these are “knockdown” experiments and there may be residual Mon1a or FCHO2 protein performing the minimal functions. Recently, FCHO1 and FCHO2 “null” cell lines were generated (73). Surprisingly, the double null cells still show endocytic coats. Unfortunately, the morphology of the Golgi apparatus and movement through the secretory pathway was not examined in this study. The study did demonstrate that the absence of FCHO2 is not lethal suggesting that its role in endocytosis and Golgi morphology maintenance are not essential or that there may be redundant pathways. To our knowledge, mammalian Mon1a “null” cells have not been generated. The FCHO2 deletion cell lines will be useful for further evaluation of the role of FCHO2 in the secretory pathway.

Most plasma membrane and secretory proteins are glycosylated and alterations in these integral macromolecules can affect their biological functions. There are human glycosylation disorders associated with defects in recycling/reorganization of the Golgi apparatus (74). That immature glycosylation moieties on proteins are present at the plasma membrane in FCHO2 or Mon1a silenced cells underscores the importance of identifying the functions these proteins in secretory pathway trafficking. What remains to be determined is their precise functions and how, when and where these proteins interact. Ongoing studies are focused on understanding these open questions.

## Author contributions

DCB performed experiments. DCB, SGM, JK and DMW analyzed data and wrote the manuscript.

## Acknowledgements

This work was supported by NIH grant HL26922 and University of Utah Funding seed grant ID:10027554 to D.M.W. D.B. was supported by NIH training grant T32DK00715. The authors express their appreciation to Joshua K. Sponbeck, Chris Rodesch, and Rishna Shrestha for technical help.

## Abbreviations

BFA: Brefeldin A
DMEM: Dulbeccos Minimal Essential Medium
DHC1: dynein-heavy chain
Dyn-IC: dynein-intermediate chain
FRAP: fluorescence recovery after photobleaching
GalNAc-T2-GFP: Golgi protein N-acetylgalactosaminyltransferase-2-green fluorescent protein
NS: nonspecific
PNS: post nuclear supernatant
VSVG-GFP: vesicular stomatitis virus G protein-green fluorescent protein.

